# RCy3: Network Biology using Cytoscape from within R

**DOI:** 10.1101/793166

**Authors:** Julia A. Gustavsen, Shraddha Pai, Ruth Isserlin, Barry Demchak, Alexander R. Pico

## Abstract

RCy3 is an R package in Bioconductor that communicates with Cytoscape via its REST API, providing access to the full feature set of Cytoscape from within the R programming environment. RCy3 has been redesigned to streamline its usage and future development as part of a broader Cytoscape Automation effort. Over 100 new functions have been added, including dozens of helper functions specifically for intuitive data overlay operations. Over 40 Cytoscape apps have implemented automation support so far, making hundreds of additional operations accessible via RCy3. Two-way conversion with networks from *igraph* and *graph* ensures interoperability with existing network biology workflows and dozens of other Bioconductor packages. These capabilities are demonstrated in a series of use cases involving public databases, enrichment analysis pipelines, shortest path algorithms and more. With RCy3, bioinformaticians will be able to quickly deliver reproducible network biology workflows as integrations of Cytoscape functions, complex custom analyses and other R packages.

## Introduction

In the domain of biology, network models serve as useful representations of interactions, whether social, neural or molecular. Since 2003, Cytoscape has provided a free, open source software platform for network analysis and visualization that has been widely adopted in biological and biomedical research fields^1^. The Cytoscape platform supports community-developed extensions, called apps, that can access third-party databases, offer new layouts, add analytical algorithms, support additional data types, and much more ^2;3^.

In 2011, the CytoscapeRPC app was created to enable R-based workflows to exercise Cytoscape v2 functionality via functions in the corresponding RCytoscape R package over XML-RPC communications protocols^4^. In 2015, the CyREST app was created to enable R-based workflows to exercise Cytoscape v3 functionality. This was achieved by the first version of the RCy3 R package, which reimplemented much of RCytoscape’s organization, data structures and syntax over REST communications protocols ^3;5^.

Here, we describe version 2.0 of the RCy3 package, which is better aligned with Cytoscape’s CyREST API. We have rewritten every function, deprecated 43 functions and added over 100 new functions. This work provides a more intuitive and productive experience for Cytoscape users learning about the RCy3 package, and it positions RCy3 to take advantage of future Cytoscape Automation ^6^ development and evolution. The goal of this paper is to describe the implementation and operation of the updated RCy3 package and to provide detailed use cases relevant to network biology applications in common and advanced bioinformatics workflows.

## Design and Implementation

RCy3 is a component of Cytoscape Automation. At the core of Cytoscape Automation is CyREST, which implements an interface to the Cytoscape Java application via the REST protocol ^6^. A collection of GET, POST, PUT and DELETE operations practically cover the complete feature set of the Cytoscape desktop software. Additional features, including those provided by user-installed Cytoscape apps, are covered by a separate command-line interface called Commands (Figure 1). All programmatic communication with Cytoscape conforms to either CyREST or Commands interfaces. For version 2.0, we redesigned the RCy3 package to parallel CyREST and Commands APIs so as to standardize the syntax and organization of its functions and streamline its future development. RCy3 functions are grouped into categories to aid the parallel development with Cytoscape Automation APIs and to facilitate navigation and comprehension of the overall package (Table 1).

**Table 1.**
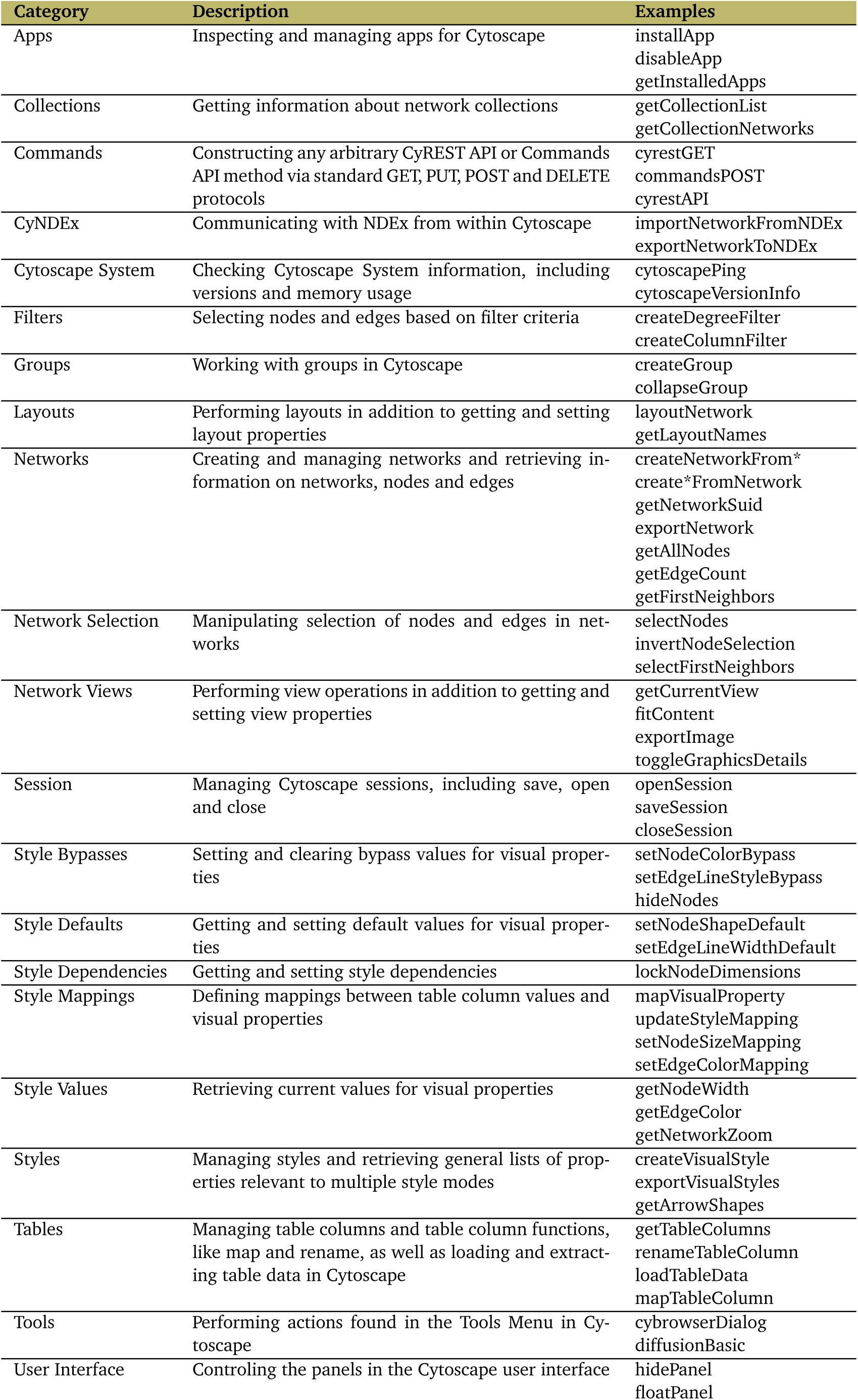
Organization of RCy3 functions. The categories correspond to separate R files in the package. Brief descriptions and example functions are listed for each category.

**Figure 1.**
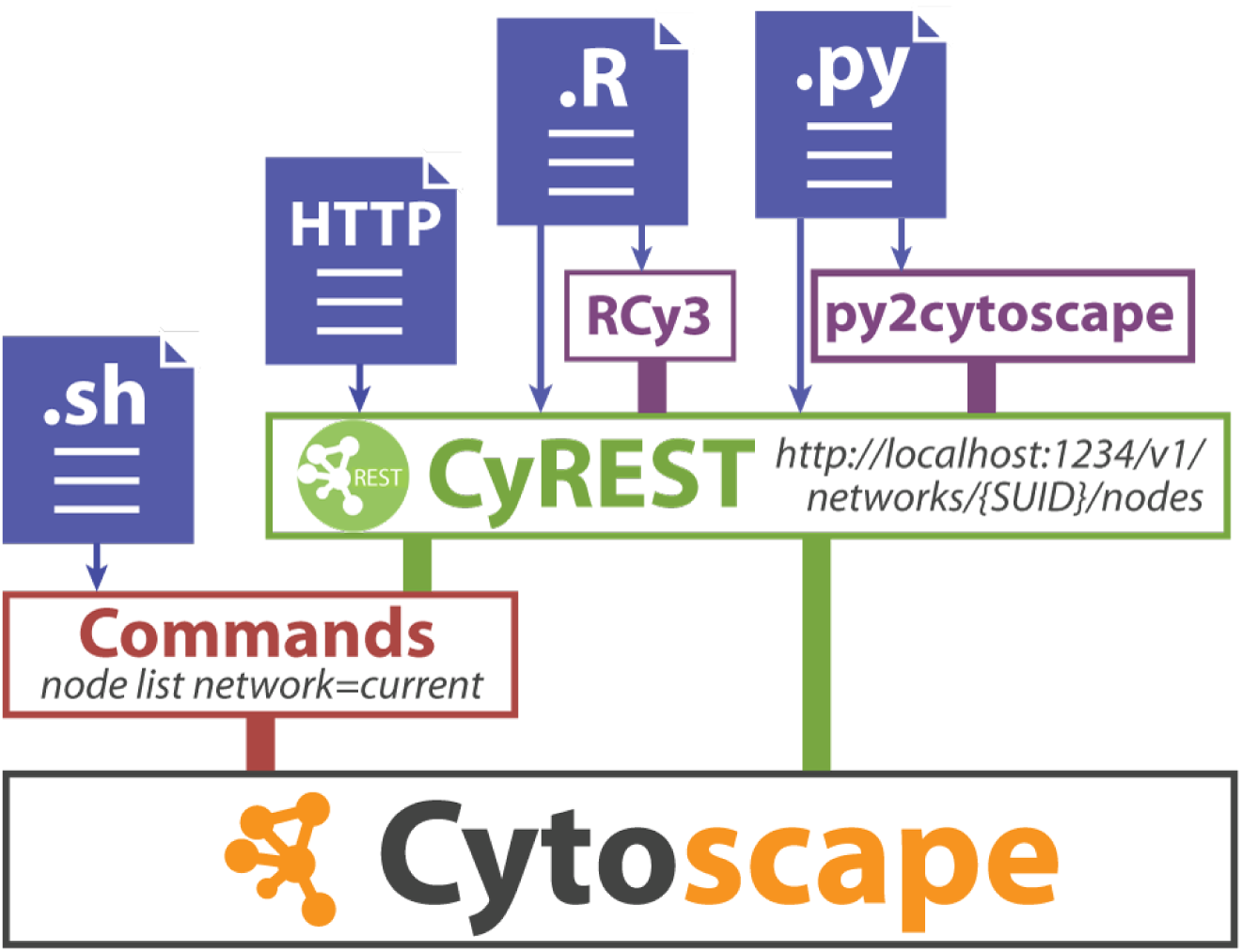
The Cytoscape Automation stack. The Java desktop version of Cytoscape (bottom) supports Commands (red) and CyREST (green) interfaces for scripting and automation either directly (blue) or via R and Python harmonization libraries (purple).

### The Basics

All RCy3 functions are ultimately implemented as calls to a small set of utility functions that execute the CyREST or Commands REST protocol (e.g., `cyrestGET` and `commandsPOST`). The internals of these functions handle the composition of operations and parameters to be sent via *httr* functions to CyREST, as well as the processing of results from JSON to appropriate R objects using *RJSONIO*.

In most RCy3 functions there is an optional argument for `base.url`. This is the URL used by RCy3 to connect to the Cytoscape desktop application via CyREST, and it defaults to ‘http://localhost:1234/v1’. The default CyREST port is 1234, and it can be changed in Cytoscape through Edit/Preferences/Properties or by command-line (see CyREST setup guide). If you change the CyREST port, you should reflect the change in the `base.url` argument per function call or change each function’s default value using the *default* package.

The second most common argument in RCy3 functions is `network.` If left as `NULL` (default), the currently selected network in the Cytoscape application is referenced. Network name or SUID (session unique identifier) can thus be explicitly specified or inferred from the current state of the application. The current network can also be controlled and retrieved by `setCurrentNetwork` and `getCurrentNetwork.` Given a `base.url` and `network` (when needed), the majority of RCy3 functions simply validate parameters and construct arguments in order to call one of the `cyrest*` or `commands*` functions.

The `commandsRun` function is a special RCy3 function that allows users to directly issue commands via Cytoscape’s command-line syntax (e.g., “node list network=current”), including commands implemented by Cytoscape app developers (see Use Cases). This single function can perform hundreds of operations made available by both Cytoscape and automation-enabled apps. Over 40 of these RCy3-supported apps are currently registered in the Cytoscape App Store ^2^. The `cyrestAPI` and `commandsAPI` open interactive Swagger documentation for the CyREST and Commands programmatic interfaces. Cytoscape Automation can be performed via these Swagger web pages. The same operations and parameters are supported by the `cyrest*` and `commands*` functions in RCy3. Command-line syntax can also be run from the Automation panel in Cytoscape, manual.cytoscape.org/en/stable/Command_Tool.html.

### Generic and Specific

The primary goal of RCy3 is to provide wrappers for every feature made available by CyREST and Commands. However, we also have a secondary goal of providing useful and intuitive functions for common workflows in R. So, in addition to the generic functions implemented to parallel the CyREST and Commands APIs, we have also implemented sets of specific helper functions.

As an example, consider the common Cytoscape operation of mapping network data values to visual style properties. CyREST has a POST endpoint for `/styles/{style name}/mappings` that takes a JSON data structure defining the mapping. We implemented `updateStyleMapping` which takes a `style.name` and mapping arguments and sends them out via `cyrestPOST.` We also implemented `mapVisualProperty` to help construct the `mapping` argument. With these generic functions one can perform any of the hundreds of visual style mappings supported by Cytoscape, including new ones added in the future. However, these functions are not simple to use, requiring knowledge of specific property names, like `“NODE_FILL_COLOR”`, and mapping data structures. To simplify usage for common situations, we therefore also implemented specific functions for over a dozen of the most commonly used mappings (e.g., `setNodeColorMapping`). With autocomplete in tools like RStudio, after just typing `setNode…` a script author is presented with a series of intuitively named functions with obvious arguments.

### Networks in R

Networks are a popular visualization option in R often implemented as graph models by *igraph* and Biocondutor’s *graph* (i.e., graphNEL). RCy3 can create networks in Cytoscape from either igraph, graphNEL or dataframe objects (`createNetworkFrom*`). Likewise, igraph and graphNEL objects can be created from networks (`create*FromNetwork`), and dataframes from node and edge tables in Cytoscape (`getTableColumns`).

In the case of `createNetworkFromDataFrames,` two dataframes are accepted as arguments, one for nodes and one for edges. The `nodes` dataframe must include a column named `“id”,` and the edges dataframe must include `“source”` and `“target”` columns. Additional columns are imported as node and edge attributes into Cytoscape. The function can also work with just one dataframe. If a dataframe of only edges is passed to `createNetworkFromDataFrames,` then a connected network will be created with all of the nodes. If a dataframe of only nodes is passed, then a network with no connections, only nodes, will be created.

RCy3 can also import network file formats supported by Cytoscape natively (e.g., SIF, xGMML and CX ^7^) and via user-installed apps (e.g., GPML ^8^ and adjacency matrices). With these functions RCy3 can interoperate with any other Bioconductor packages that deal with networks in a standardized manner, providing advanced network visualization options and advanced network analytics from the Cytoscape ecosystem (see Table 2).

**Table 2.**
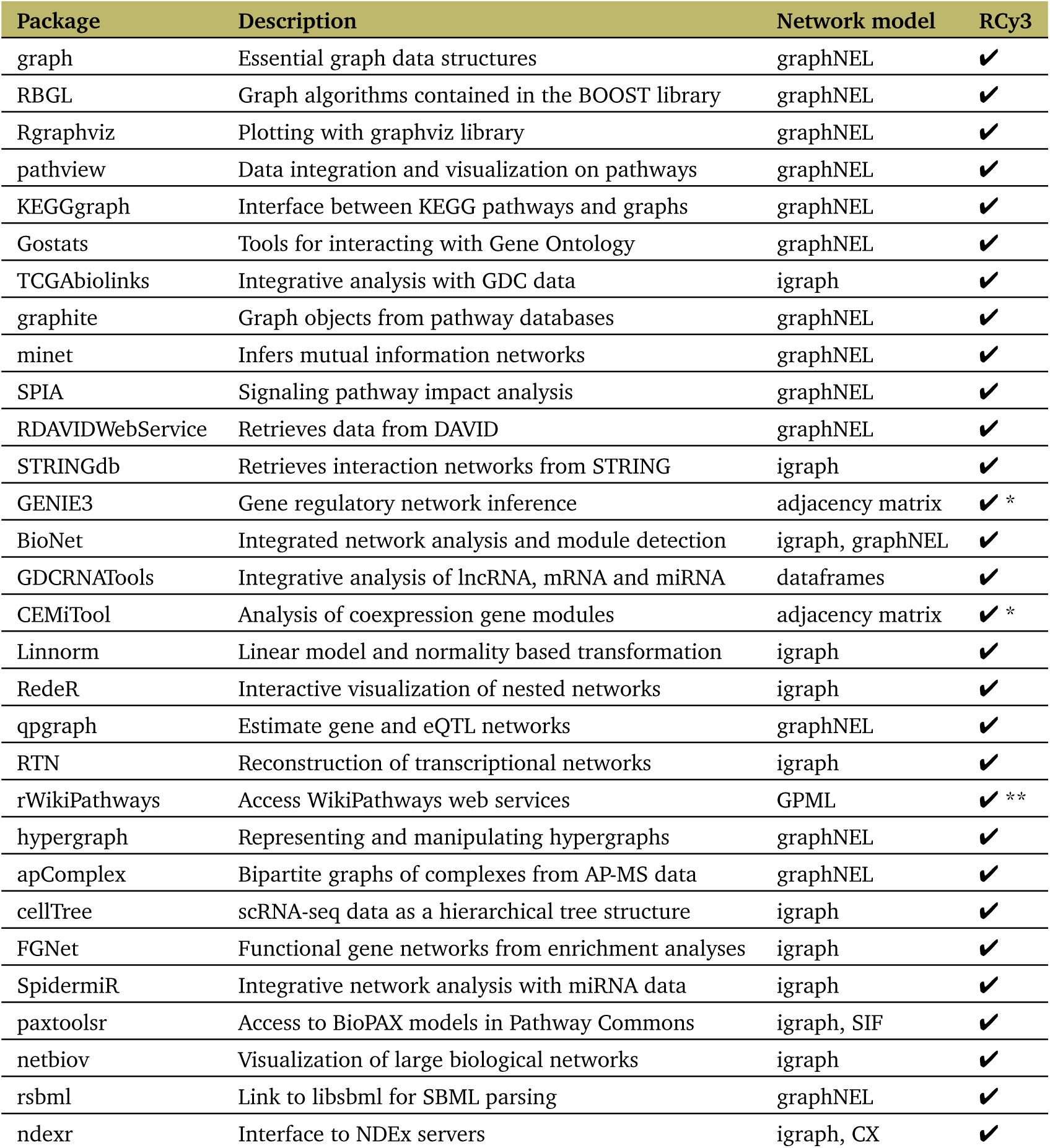
Top 30 network-related Bioconductor packages ordered by rank (as of Sept 2019) and their network models relevant to RCy3. RCy3 can provide and consume network models to and from these packages (checkmarks). Asterisks indicate where a Cytoscape app is required (* = aMatReader, ** = WikiPathways).

## Operation

In order to work with RCy3 you must have Cytoscape v3.7 or later installed and running. Cytoscape can be installed from cytoscape.org. The RCy3 package can be installed from Bioconductor:

~~~
if (!requireNamespace(“BiocManager”, quietly = TRUE))
    install.packages(“BiocManager”)
BiocManager∷install(“RCy3”)
library(RCy3)
~~~

Launch Cytoscape and keep it running whenever using RCy3. Confirm that you have everything installed and that RCy3 is communicating with Cytoscape via CyREST:

~~~
cytoscapePing ()
#[1] “You are connected to Cytoscape!”
~~~

As with any R package, one can access the documentation and browse over a dozen vignettes included in the RCy3 package:

~~~
help(package=RCy3)
browseVignettes(“RCy3”)
~~~

## Use Cases

The following sections demonstrate a variety of common and advanced network biology use cases as runnable R code snippets. The code for these use cases is also available as an online Rmd notebook and Rmd file in the Cytoscape Automation repository (see Data Availability). The first set focuses on fundamental Cytoscape operations that are common to most use cases:

- Loading networks (from R objects, Cytoscape files and public databases)
- Visualizing network data
- Filtering by node degree or data
- Saving and exporting networks

Additionally, there are examples that demonstrate analytical workflows, relying not only on Cytoscape, but also Cytoscape apps and other R packages:

- Building maps of enrichment analysis results using EnrichmentMap and AutoAnnotate
- Visualizing integrated network analysis using BioNet
- Performing advanced graph analytics using RBGL

### Loading Networks

Networks come in all shapes and sizes, in multiple formats from multiple sources. The following code snippets demonstrate just a few of the myriad ways to load networks into Cytoscape using RCy3.

From R objects…

~~~
# From graph objects (graphNEL)
g <- makeSimpleGraph()
createNetworkFromGraph(g)
## And round-trip back from Cytoscape to graph
g2 <- createGraphFromNetwork()

# From igraph objects library(igraph)
ig <- igraph∷make_graph(“Zachary”)
createNetworkFromIgraph(ig)
## And round-trip back from Cytoscape to igraph
ig2 <- createIgraphFromNetwork()
## Note that the Cytoscape model infers directionality

# From dataframes
nodes <- data.frame(id=c(“node 0”,”node 1”,”node 2”,”node 3”),
                          group=c(“A”,”A”,”B”,”B”), #categorical strings
                          score=as.integer(c(20,10,15,5)), #integers
                          stringsAsFactors=FALSE)
edges <- data.frame(source=c(“node 0”,”node 0”,”node 0”,”node 2”),
                          target=c(“node 1”,”node 2”,”node 3”,”node 3”),
                          interaction=c(“inhibits”,”interacts”,
                                        “activates”,”interacts”), #optional
                          weight=c(5.1,3.0,5.2,9.9), #numerics
                          stringsAsFactors=FALSE)
createNetworkFromDataFrames(nodes, edges)
~~~

From Cytoscape-supported file formats…

~~~
# From Cytoscape session files
## Will erase and replace all data from current session!
openSession() # default file = galFiltered.cys

# From local network files
importNetworkFromFile() # default file = galFiltered.sif
## Supported file formats: SIF, GML, xGMML, graphML, CX, plus

# From NDEx (https://ndexbio.org), the network database
importNetworkFromNDEx(“5be85817-1e5f-11e8-b939-0ac135e8bacf”)
## Account information or accessKey are required arguments only
## when accessing private content
~~~

From public databases via Cytoscape apps…

~~~
# From STRING (https://string-db.org), starting with a list of genes/proteins
installApp(“stringApp”) # http://apps.cytoscape.org/apps/stringapp
gene.list <- c(“T53”,”AKT1”,”CDKN1A”)
gene.str <- paste(gene.list, collapse = “,”)
string.cmd <- paste(“string protein query cutoff=0.99 limit=40 query”,
                           gene.str, sep = “=“)
commandsRun(string.cmd)

# From WikiPathways (https://wikipathways.org), starting with a keyword
library(rWikiPathways) # install from Bioconductor
installApp(“WikiPathways”) # http://apps.cytoscape.org/apps/wikipathways
keyword <- “glioblastoma”
gbm.pathways <- findPathwaysByText(keyword)
gbm.wpid <- gbm.pathways[[1]]$id # let’s just take the first one
wikipathways.cmd <- paste(“wikipathways import-as-pathway id”,
                                 gbm.wpid, sep = “=“)
commandsRun(wikipathways.cmd)
~~~

From public databases via Cytoscape apps…

### Visualizing Data on Networks

Cytoscape excels at generating publication-quality network visualization with data overlays. This vignette demonstrates just one of the hundreds of visual style mapping options using RCy3.

~~~
# Load sample network
closeSession(FALSE) # clears all session data wihtout saving
importNetworkFromFile() # default file = galFiltered.sif

# Load sample data
csv <- system.file(“extdata”,”galExpData.csv”, package=“RCy3”)
data <- read.csv(csv, stringsAsFactors = FALSE)
loadTableData(data,data.key.column=“name”)

# Prepare data-mapping points
gal80Rexp.min <- min(data$gal80Rexp, na.rm = T)
gal80Rexp.max <- max(data$gal80Rexp, na.rm = T)
## For a balanced color gradient, pick the largest absolute value
gal80Rexp.max.abs <- max(abs(gal80Rexp.min), abs(gal80Rexp.max))

# Set node color gradient from blue to white to red
setNodeColorMapping(‘gal80Rexp’, c(-gal80Rexp.max.abs, 0, gal80Rexp.max.abs),
                    c(‘#5577FF’,’#FFFFFF’,’#FF7755’), default.color = ‘#999999’)
~~~

### Filtering Networks by Degree and by Data

Network topology and associated node or edge data can be used to make selections in Cytoscape that enable filtering and subnetworking. The filters are added to the Select tab in the Control Panel of Cytoscape’s GUI and saved in session files.

~~~
# Load demo Cytoscape session file
openSession() # default file = galFiltered.cys
net.suid <- getNetworkSuid() # get SUID for future reference

# Filter for neighbors of high degree nodes
createDegreeFilter(filter.name = “degree filter”,
                   criterion = c(0,9),
                   predicate = “IS_NOT_BETWEEN”)
selectFirstNeighbors() # expand selection to first neighbors
createSubnetwork(subnetwork.name = “first neighbors of high degree nodes”)

# Filter for high edge betweenness
createColumnFilter(filter.name = “edge betweenness”,
                   type = “edges”,
                   column = “EdgeBetweenness”,
                   4000,
                   “GREATER_THAN”,
                   network = net.suid)
createSubnetwork(subnetwork.name = “high edge betweenness”)
~~~

### Saving and Exporting Networks

There are local and cloud-hosted options for saving and sharing network models and images. The Cytoscape session file (CYS) includes all networks, collections, tables and styles. It retains every aspect of your session, including the size of the application window. Network and image exports include only the currently active network. Export to NDEx requires account information you can obtain from ndexbio.org. Files are saved to the current working directory by default, unless a full path is provided.

~~~
# Saving sessions
saveSession(“MySession”) #.cys
## Leave filename blank to update previously saved session file

# Exporting images and networks
exportNetwork() #.sif
## Optionally specify filename, default is network name
## Optionally specify type: SIF(default), CX, cyjs, graphML, NNF, SIF, xGMML
exportImage() #.png
## Optionally specify filename, default is network name
## Optionally specify type: PNG (default), JPEG, PDF, PostScript, SVG

# Exporting to NDEx, a.k.a. “Dropbox” for networks
exportNetworkToNDEx(username, password, TRUE)
## Account information (username and password) is required to upload
## Use updateNetworkInNDEx if the network has previously been uploaded
~~~

### Building Maps of Enrichment Analysis Results

This workflow illustrates how to plot an annotated map of enrichment results using the EnrichmentMap Pipeline Collection of apps in Cytoscape ^9^. An enrichment map is a network visualization of related genesets in which nodes are gene sets (or pathways) and edge weight indicates the overlap in member genes ^10^. Following the construction of the enrichment map, AutoAnnotate clusters redundant gene sets and uses WordCloud ^11^ to label the resulting cluster (Figure 2). The code uses the Commands interface to invoke EnrichmentMap and AutoAnnotate apps. After installing apps, run commandsAPI() to open the live Swagger documentation to browse and execute command-line syntax.

**Figure 2.**
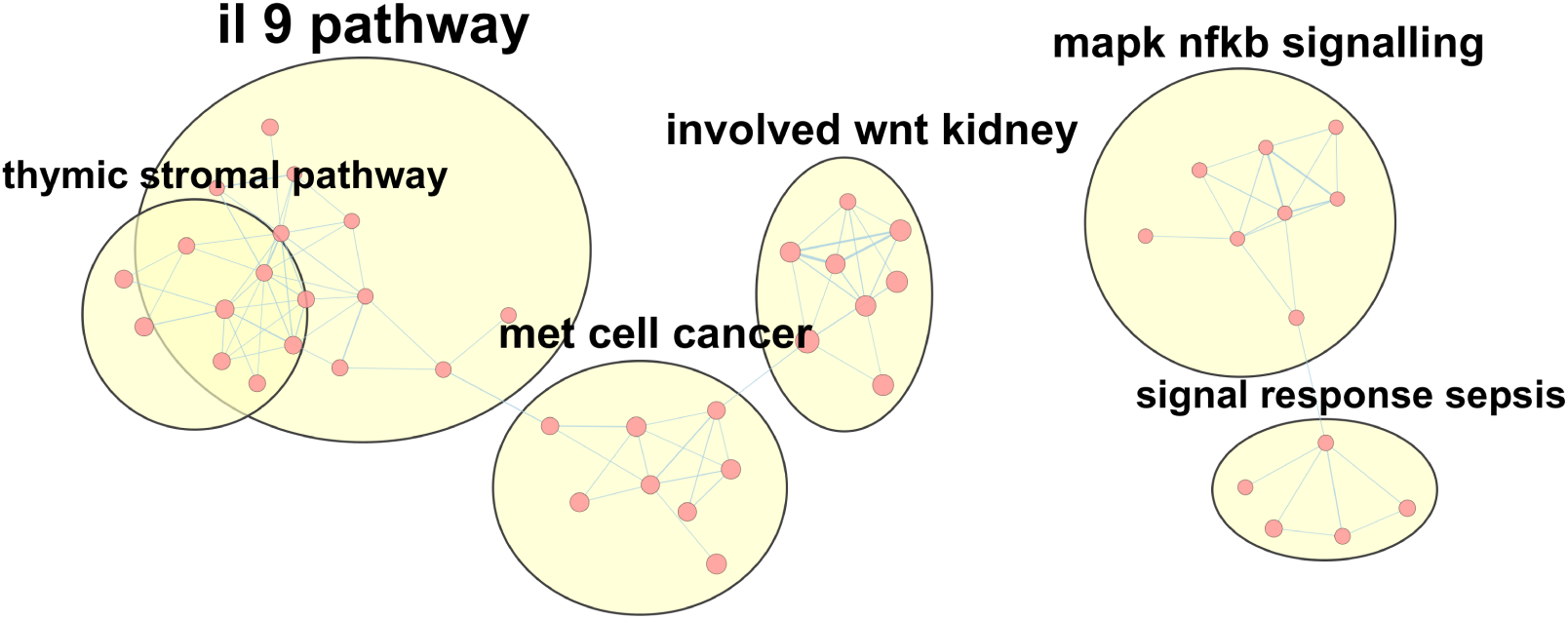
Annotated enrichment map of pathways. A few of the largest clusters of pathways from WikiPathways are displayed as a network with WordCloud-based labels annotating partially overlapping groups (yellow areas). The size of the labels correspond to the size of the groups.

~~~
installApp(“EnrichmentMap Pipeline Collection”) # installs 4 apps

# Optionally browse commands for enrichmentmap
commandsAPI()

# Download sample gmt file of human pathways
gmt.file <- “rcy3_enrichmentmap.gmt”
download.file(file.path(“http://download.baderlab.org/EM_Genesets”,
                        “September_01_2019/Human/symbol/Pathways”,
                        “Human_WikiPathways_September_01_2019_symbol.gmt”),
              gmt.file)

# Run EnrichmentMap build command
em_command <- paste(‘enrichmentmap build analysisType=“generic”’,
                           “gmtFile=“, paste(getwd(), gmt.file, sep=“/”),
                           “pvalue=“, 1,
                           “qvalue=“, 1,
                           “similaritycutoff=“,0.25,
                           “coefficients=“,”JACCARD”)
print(em_command)
commandsGET(em_command)

# Run the AutoAnnotate command
aa_command <- paste(“autoannotate annotate-clusterBoosted”,
                           “clusterAlgorithm=MCL”,
                           “labelColumn=EnrichmentMap∷GS_DESCR”,
                           “maxWords=3”)
print(aa_command)
commandsGET(aa_command)

# Annotate a subnetwork
createSubnetwork(c(1:4),” mclCluster”)
commandsGET(aa_command)
~~~

### Visualizing Integrated Network Analysis Using *BioNet*

The *BioNet* package implements analytical methods to perform integrated network analysis, for example, of gene expression data and clinical survival data in the context of protein-protein interaction networks. Partnered with RCy3, the analytical results from *BioNet* can be visualized in Cytoscape with vastly more options for customization. Starting with the “Quick Start” tutorial from *BioNet*, we pass the results directly to Cytoscape for visualization:

~~~
library(BioNet) # install from Bioconductor
library(DLBCL) # install from Bioconductor
data(dataLym)
data(interactome)

## The following steps are from BioNet*’*s Quick Start tutorial:
pvals <- cbind(t = dataLym$t.pval, s = dataLym$s.pval)
rownames(pvals) <- dataLym$label
pval <- aggrPvals(pvals, order = 2, plot = FALSE)
subnet <- subNetwork(dataLym$label, interactome)
subnet <- rmSelfLoops(subnet)
fb <- fitBumModel(pval, plot = FALSE)
scores <- scoreNodes(subnet, fb, fdr = 0.001)
module <- runFastHeinz(subnet, scores)
logFC <- dataLym$diff
names(logFC) <- dataLym$label
plotModule(module, scores = scores, diff.expr = logFC)

# Using RCy3 we can generate a custom visualization of the same output
## Create network from graphNEL object and load data calculated above
createNetworkFromGraph(module, “module”, “BioNet”)
loadTableData(as.data.frame(scores))
loadTableData(as.data.frame(logFC))

## Set styles
setNodeLabelMapping(“geneSymbol”)
setNodeFontSizeDefault(18)
setNodeBorderWidthDefault(3.0)
logFC.max.abs <- max(abs(min(logFC)), abs(max(logFC)))
setNodeColorMapping(‘logFC’, c(-logFC.max.abs, 0, logFC.max.abs),
                    c(‘#5577FF’,’#FFFFFF’,’#FF7755’), default.color = ‘#999999’)
createColumnFilter(“Positive scores”, “scores”,c(0,max(scores)),”BETWEEN”)
setNodeShapeBypass(getSelectedNodes(), “ELLIPSE”)
~~~

### Performing Advanced Graph Analytics Using *RBGL*

As an interface to the BOOST library, the *RBGL* Bioconductor package offers an impressive array of analytical functions for graphs. Here we will start with a network in Cytoscape, load it into R as a *graph* object, perform shortest path calculation using *RBGL* and then visualize the results back in Cytoscape (Figure 3).

**Figure 3.**
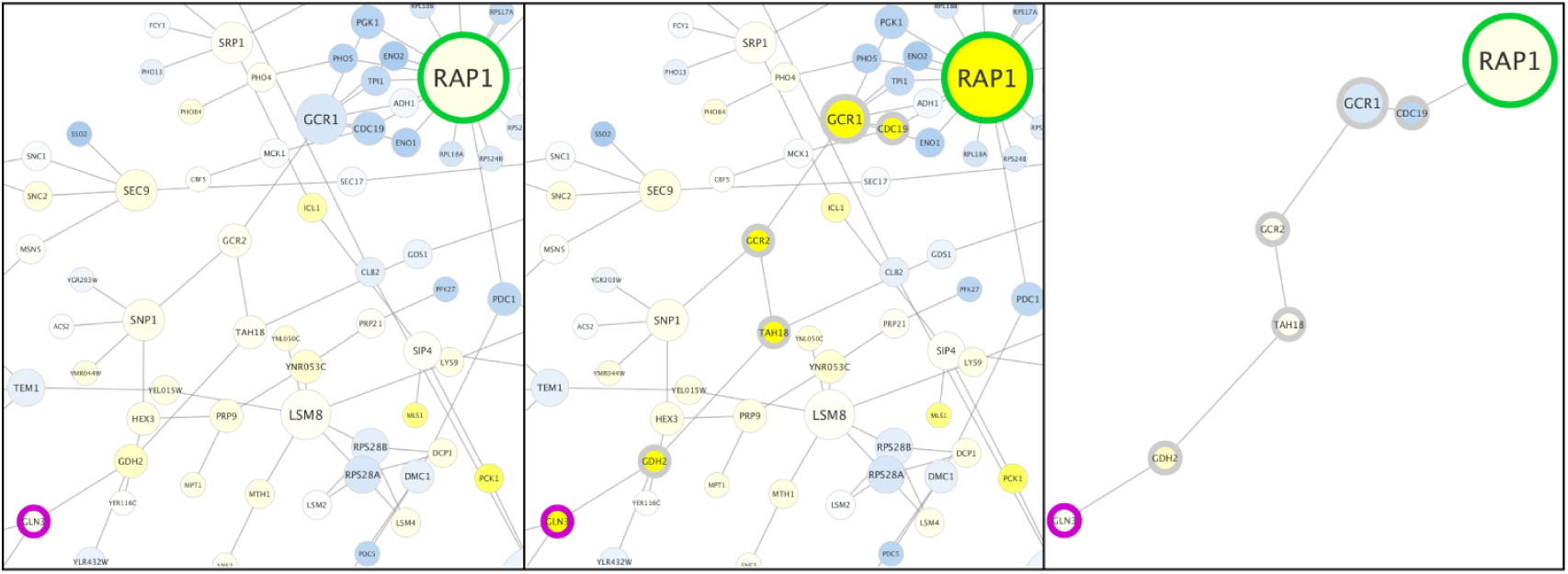
Visualizing shortest path results. In Cytoscape, the start and finish nodes were colored green and magenta, respectively (left panel). The shortest path calculated by *RBGL* was then selected and nodes along the path were highlighted with thick borders (middle panel). Finally, a subnetwork was created from the selected path (right panel). Note that while common gene names are displayed as node labels, official yeast identifiers are the actual node names that are referenced in the script.

~~~
library(RBGL) # install from Bioconductor
# Convert a sample Cytoscape network to a graph object
openSession()
g <- createGraphFromNetwork()

# Identify start and finish nodes (styling is optional)
start <- “YNL216W”
finish <- “YER040W”
setNodeBorderWidthBypass(c(start,finish), 20)
setNodeBorderColorBypass(start,”#00CC33”)
setNodeBorderColorBypass(finish,”#CC00CC”)

# Use RBGL to perform shortest path calculation
shortest <- sp.between(g, start, finish)
shortest$’YNL216W:YER040W’$length
#[1] 6
shortest.path <- shortest$’YNL216W:YER040W’$path_detail

# Visualize results in Cytoscape
selectNodes(shortest.path, “name”)
setNodeBorderWidthBypass(shortest.path, 20)
createSubnetwork()
~~~

## Discussion

Every operation exposed by Cytoscape’s REST API has now been implemented as a function in RCy3 2.0. Furthermore, RCy3 provides dozens of higher-level helper functions in support of common usage, such as `setNodeColorMapping,` to make script writing more intuitive and efficient. The issue trackers for CyREST and RCy3 are linked for functions pending implementation, such as `mergeNetworks,` as well as for bug fixes. Thus, RCy3 is expected to keep pace with future development of Cytoscape Automation. More broadly, RCy3 is an integral part of the Cytoscape ecosystem, which includes the network repository NDEx ^7^, Cytoscape apps and services, and web components like cytoscape.js _?_. The ecosystem has also defined interfaces and standard formats to interoperate with interaction databases and annotation services. Adopting RCy3 for network analysis will establish a connection to the Cytoscape ecosystem and enable Cytoscape Automation workflows _6_. As the sharing and publishing of analysis scripts and workflows become more routine (if not mandated), software tools designed to work in an open and accessible ecosystem are becoming essential.

## Data availability

- RCy3 vignettes: https://bioconductor.org/packages/release/bioc/html/RCy3.html
- RCy3 Rmd notebooks: https://cytoscape.org/cytoscape-automation/for-scripters/R/notebooks/
- RCy3 workshop presentations: http://tutorials.cytoscape.org/#automation
- Video demonstrations of RCy3: https://www.youtube.com/playlist
- Cytoscape Automation training: http://automation.cytoscape.org/
- Cytoscape Automation code repository: https://github.com/cytoscape/cytoscape-automation

## Software availability

- Bioconductor: https://bioconductor.org/packages/release/bioc/html/RCy3.html
- GitHub: https://github.com/cytoscape/RCy3
- Issue tracker: https://github.com/cytoscape/RCy3/issues

## Author contributions

AP, SP and RI redesigned and implemented version 2 of RCy3. AP, JAG, SP and BD drafted the manuscript. AP, JAG, SP and RI contributed use cases.

## Acknowledgments

We would like to acknowledge contributions by other developers on the original implementation of RCy3 by Paul Shannon, Tanja Muetze and Georgi Kolishkovski. We also greatly appreciate the input from Mark Grimes (testing), Martin Morgan (Bioconductor), and the excellent work on CyREST and Cytoscape Commands by David Otasek and John “Scooter” Morris.

